# Immunoglobulin profile and B-cell frequencies are altered with changes in the cellular micro-environment independent of the stimulation conditions

**DOI:** 10.1101/818153

**Authors:** Dannielle K Moore, Gina R Leisching, Candice I Snyders, Andrea Gutschmidt, Ilana C van Rensburg, Andre G Loxton, the SU-IRG Consortium

**Author notes:** Andre G Loxton: (Corresponding author). Summary Statement: B-cell behaviour is dependent on the presence of other immunogenic cells.

## Abstract

B-cells are essential in the defense against *Mycobacterium tuberculosis*. Studies on isolated cells may not accurately reflect the responses that occur *in vivo* due to the presence of other cells. This study elucidated the influence of microenvironment complexity on B-cell polarisation and function in the context of TB disease. B-cell function was tested in whole blood, PBMC’s and as isolated cells. The different fractions were stimulated and the B-cell phenotype and immunoglobulin profiles analysed. The immunoglobulin profile and killer B-cell frequencies varied for each of the investigated sample types, while in an isolated cellular environment, secretion of immunoglobulin isotypes IgA, IgG2 and IgG3 was hampered. The differences in the immunoglobulin profile highlight the importance of cell-cell communication for B-cell activation. In contrast, increased frequencies of killer B-cells were observed following cellular isolation, suggesting a biased shift in augmented immune response *in vitro*. This suggests that humoral B-cell function and development was impaired likely due to a lack of co-stimulatory signals from other cell types. Thus, B-cell function should ideally be studied in a PBMC or whole blood fraction.

## Introduction

Over the past two decades, researchers have increased the use of cell culture and isolated tissue samples for biological research as an alternative to *in vivo* animal studies, due to the large cost and strict regulatory conditions involved (Murphy, 1991; Brown Jr, 1997; Adams and Larson, 2007). For this reason, cell culture studies have formed the fundamental basis of a variety of research topics (4). While the information gained from these isolated cell studies provides valuable insight into biological mechanisms under investigation, they do not account for the many factors that control these physiological responses *in vivo*. Numerous studies have illustrated the ability of various cell types to modulate the host immune response under different conditions (Blair *et al.*, 2010b; Carter, Rosser and Mauri, 2012a), and the presence and activation of these cell types may contribute significantly to the function of a cell type of interest, through directing the mounting immune response. As such, absence of these cells during isolated cell studies may result in artificial observations and inaccurate assumptions regarding the role of a cell population during health and disease. The effects of isolation on investigated cellular responses is evident in many studies (Kondo and Magee, 1977; Sanders *et al.*, 1983; Murphy, 1991), in which a particular condition produced a measured immune response in whole organism, while having minimal or no effect on isolated target cells or tissue (vice versa).

Recent studies investigating the role of B-cells during tuberculosis (TB) revealed impaired B-cell function and decreased B-cell frequencies during active disease (Joosten *et al.*, 2016a; Van Rensburg *et al.*, 2017). A regulatory B-cell (B_reg_) subtype, killer B-cells, was recently discovered and has been implicated in a variety of immune conditions, including M.*tb* infection (Lundy et al., 2015; van Rensburg and Loxton, 2018a). Emerging evidence has acknowledged B-cells as essential in the defense against *Mycobacterium tuberculosis (M.tb)*. Prior to the participation of B-cells in host immune responses, development and migration of precursor cells known as transitional B-cells from the bone marrow to the spleen is required; where they give rise to mature B-cells in response to antigenic stimulation (Paul, 2013; Abbas, Lichtman and Pillai, 2014). Immature transitional 1 (T1) B-cells (CD19^+^CD21^−^CD23^−^) form the foundation from which transitional 2 (T2) B-cells (CD19^+^CD21^+^CD23^+^) and mature B-cells sequentially derive (Petro *et al.*, 2002). However, T2 B-cells have been found to be more receptive to cellular activation and proliferation, in comparison to T1 B-cells (Mackay and Browning, 2002; Chung, Silverman and Monroe, 2003). As such, the presence and regulation of these T2 B-cells dramatically affects the course of the mounted immune responses.

Functional analysis of these cells in numerous health settings identified the role of these cells as potential immune modulators responsible for controlling the immune response during disease and infection. (Lundy and Boros, 2002; Matsushita *et al.*, 2008; Lundy, 2009; Chesneau *et al.*, 2013; Lundy, Klinker and Fox, 2015). A study by van Rensburg and colleagues (van Rensburg *et al.*, 2017) investigated the frequency of these killer B_regs_ during active TB disease and found a decrease in the frequency of these cells during TB diseased individuals when compared to controls. Upon successful TB treatment, these killer B-cell frequencies returned to levels comparable to that of healthy controls, suggesting a pivotal role of these B_regs_ in protective anti-TB immunity (van Rensburg *et al.*, 2016; Van Rensburg *et al.*, 2017).

The objective of this study was to elucidate the influence of microenvironment complexity on B-cell polarisation and function in the context of TB disease. Current research findings inferring the role of B-cells during *M.tb* infection utilized isolated B-cell cultures to investigate their functional capacity. Thus we sought to firstly assess and characterize B-cell function in whole blood, PBMCs and as an isolated culture in response to various antigenic stimuli, and secondly to determine the significance of the use of isolated cell culture techniques in studies inferring the role of B-cells during TB disease.

## Results

### The B-cell immunoglobulin profile varies considerably according to cellular microenvironment/sample fraction

The mechanisms employed by B-cells toward facilitating enhanced anti-TB immunity remains unresolved. Subsequently, the effects of microenvironment complexity i.e. the degree of cellular isolation on B-cell function was investigated. This was achieved by examining the immunoglobulin expression patterns within culture supernatants following stimulation of B-cells in various sample fractions (whole blood, PBMCs and isolated B-cells).

Altered immunoglobulin profiles were observed across the various sample fractions, regardless of stimulation conditions, however most notable was the observation of increased IgM release from isolated B-cells (Figure 1). A significant difference (p<0.05) in the immunoglobulin profile was observed for all isotypes, except IgG4, following isolation of B-cells from whole blood (Figure 2a). In some instances, the relative abundance of an isotype within a given sample set, such as IgE (p=0.0001), IgG3 (p=0.0000) and IgA (ns, p>0.05), was observed to increase following PBMC isolation. However, in majority of cases a decrease in the relative abundance of an immunoglobulin isotype, specifically IgG1 (p = 0.0000), IgG2 (ns, p>0.05) IgG4 (ns, p>0.05) and IgM (ns, p>0.05), within a given sample set was observed following PBMC isolation. A significant decrease in the relative abundance of most isotypes, including IgA (p=0.0000), IgG1 (p=0.0000) and IgG2 (p=0.0000), was observed following isolation of B-cells compared to whole blood, whilst an increase in the relative abundance of IgE (p=0.0004) and IgM (p=0.0000) was found. Likewise, a substantial decrease in the relative abundance of most isotypes, IgA (p=0.0000), IgG1(p=0.0001), IgG2 (p=0.0000) and IgG3 (p=0.0000), was observed following isolation of B-cells compared to the PBMC sample fraction, whereas an increase in the relative abundance of IgM (p=0.0000) was found. Incidentally, the effect of sample type on the abundance of the various immunoglobulin isotypes was investigated to discern the effect of cellular isolation on the magnitude of subsequent B-cell responses. A significant difference in quantified immunoglobulin levels was observed for all isotypes following each successive isolation procedure (data not shown). To account for the limitations involved in comparing plasma supernatants to culture supernatants, only differences between PBMCs and isolated B-cells were considered. Here, significant decreases in the observed concentration of all immunoglobulin isotypes were observed for isolated B-cell culture supernatants compared to those of PBMC samples.

**Figure 1.**
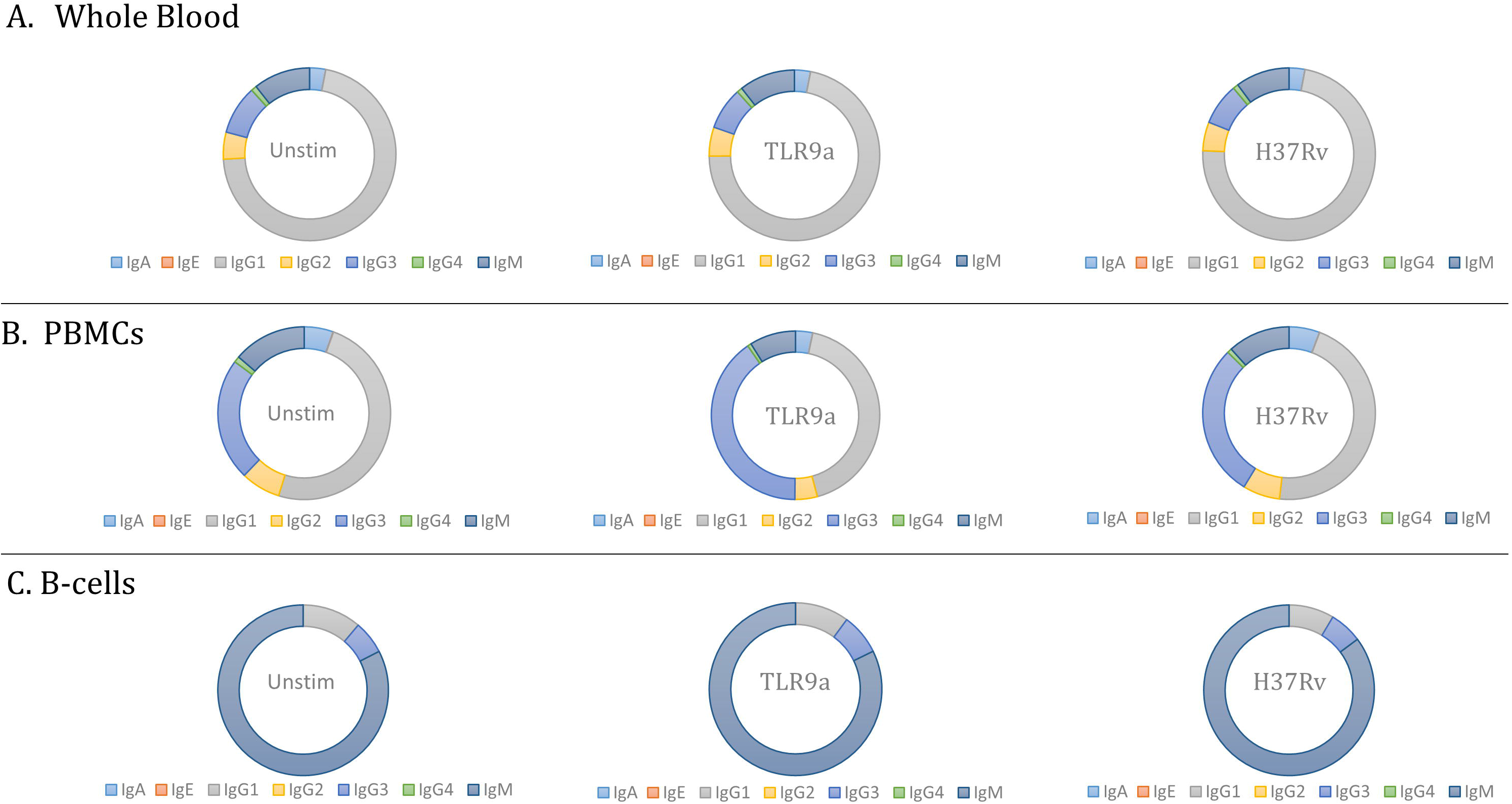
Immunoglobulin profile of supernatants obtained from each of the stimulatory conditions of the various cellular fractions. Following a 24-hour stimulatory period, plasma/culture supernatants of each of the stimulatory conditions for all sample types was collected and the immunoglobulin secretion profile determined using Luminex. (a) Representation of the average secretion of each isotype within whole blood for each stimulatory condition (b) Representation of the average secretion of each isotype within the PBMC fraction for each stimulatory condition (c) Representation of the average secretion of each isotype within the isolated B-cell fraction for each stimulatory condition, n=23.

**Figure 2.**
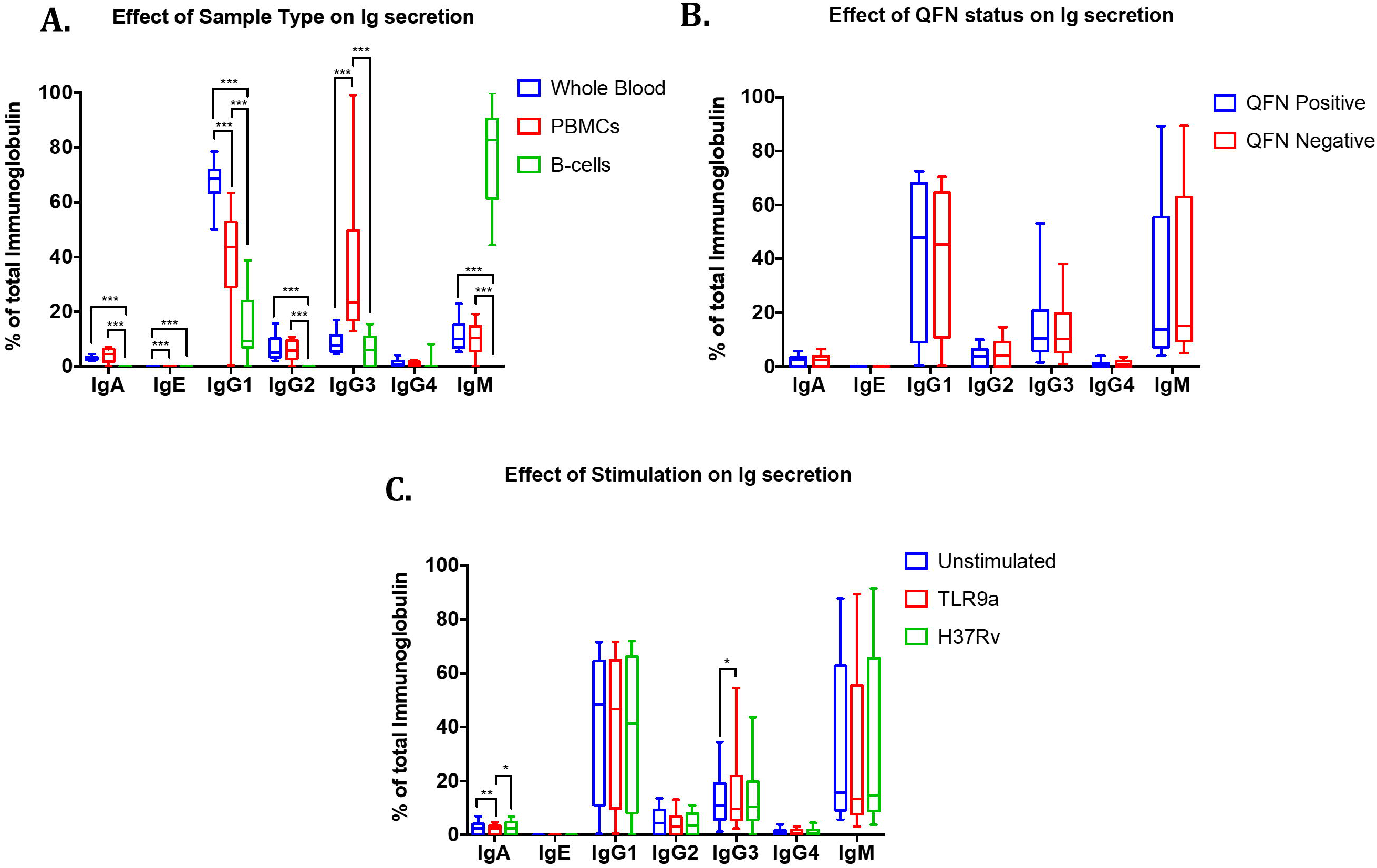
Analysis of developmental B-cell phenotypic frequencies. Following a 24-hour stimulatory period, cells from each of the stimulatory conditions, for all sample types, was collected and the B-cell phenotypic frequencies determined using flow cytometry. Notable, statistical significance was only investigated for whole blood and PBMC sample type as isolated B-cell samples for the respective QFN groups were pooled (according to stimulation condition) prior to flow analysis, due to inadequate cell numbers. Whiskers denote 10-90 percentile. Statistical differences between culture conditions was calculated using a four-way mixed model ANOVA. Comparisons within groups was calculated using the Fishers LSD post-hoc test. A two-way step-up Benjamini, Krieger and Yekutieli False Discovery rate (FDR) approach, with a FDR of 1%, was used to correct for multiple testing. Statistical significance is indicated by an asterisk, in which the p < 0.05 (*), p<. 0.01 (**) and p<0.001(***). (a) Effects of sample type on the developmental state of B-cells (b) Effects of Quantiferon (QFN) status on the developmental state of B-cells (c) Effects of stimulation condition on the developmental state of B-cells.

Additionally, the effects of QFN status and stimulation condition on immunoglobulin profiles were investigated. We investigated whether or not *M.tb*-exposed (QFN positive) individuals with immune memory would respond differently to *M.tb* challenge compared to QFN negative individuals. In all instances, QFN status was found to have no effect on the relative abundance of the various immunoglobulin isotypes (Fig. 2B). To conclude, the impact of stimulation condition on immunoglobulin profiles were investigated to determine the effect of *M.tb* infection on B-cell performance. For the majority of immunoglobulin isotypes, stimulation with different antigens had no effect on the measured immunoglobulin abundance (Fig. 2C). However, a significant decrease in IgA levels were observed following TLR9 stimulation for all sample types compared to unstimulated controls (p=0.0038) and H37Rv stimulated (p=0.0292) samples. Conversely, an increase in IgG3 levels were observed following TLR9 stimulation for all sample types compared to unstimulated controls (p=0.0482).

### B-cells isolation results in decreased frequencies of MZ B-cells

In addition to investigating alterations within the immunoglobulin profile of B-cells, variations in the phenotypic frequencies of various B-cell subsets was investigated to evaluate the effect of microenvironment complexity on cell function. The expression of the cell surface receptors CD21 and CD23 by B-cells was examined to determine the proportion of B-cells within various developmental stages following antigenic stimulation. Following a 24-hour stimulatory period, cells from all stimulatory conditions for each of the cellular fractions was collected and phenotypic frequencies of the various B-cell population determined using flow cytometry. It is important to note that due to limited cell numbers, isolated B-cell samples (B-cells only) were pooled according to QFN status and stimulatory conditions prior to flow cytometry analysis, prohibiting the assessment of individual sample distribution and statistically significant differences between B-cells only and other microenvironment conditions. Consequently, observations made from the resulting data focus principally on the difference between whole blood and PBMC, while inferring the physiological implications of the trends observed for isolated B-cells. The effect of sample type on B-cell development was investigated, in which no significant difference in T1, T2, MZ and FO B-cell frequencies was observed between whole blood and PBMCs. However, a shared pattern of decreased CD19^+^CD21^+^CD23^−^ (MZ) B-cells, whilst an increase in CD19^+^CD21^+^CD23^+^ (T2) B-cells, CD19^+^CD21^−^CD23^+^ (FO) B-cells and CD19^+^CD21^−^CD23^−^ (T1) B-cells was observed for all sample types following each successive isolation procedure (Fig. 3A). Regrettably, the significance of alterations in these B-cell frequencies for isolated B-cell samples cannot be analyzed, however a sizeable difference in the investigated frequencies is apparent. These results indicate the potential impact that B-cell isolation has on maturation in response to stimulation *in vitro*.

**Figure 3.**
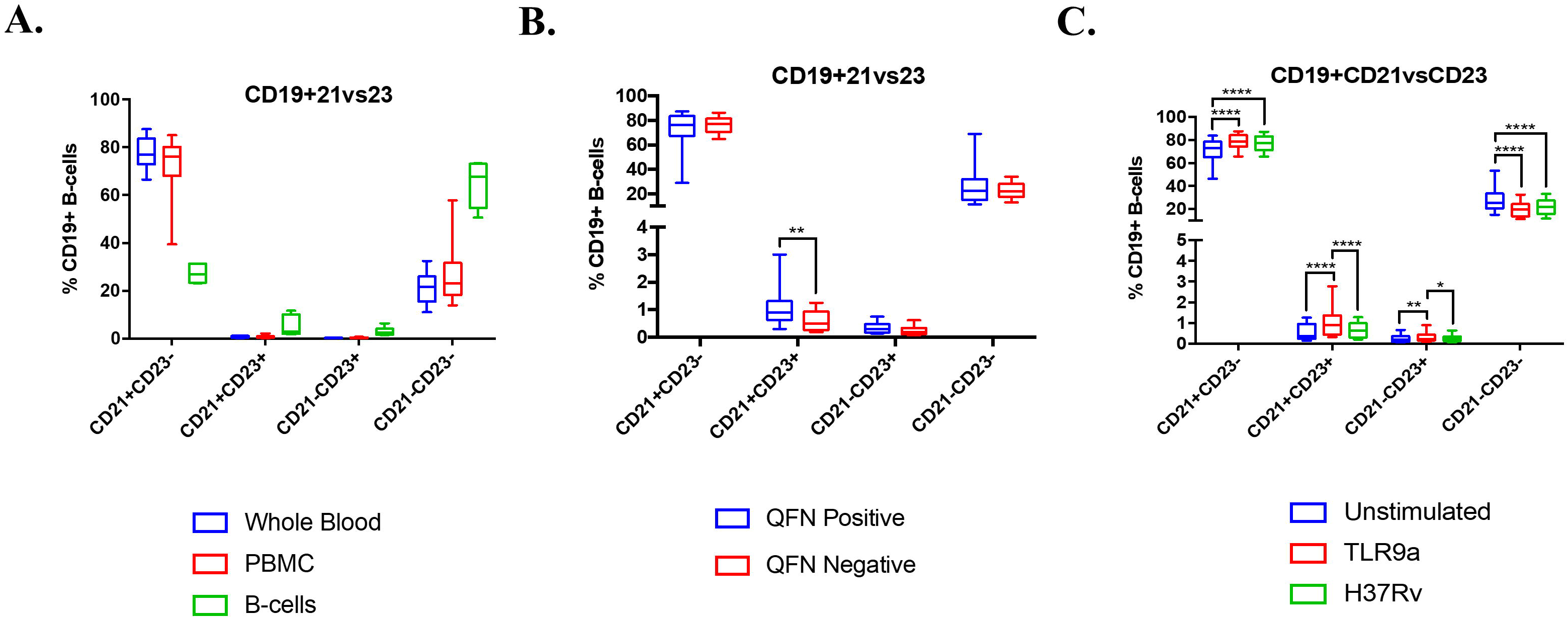
Evaluation of the effect of various experimental factors on B-cell immunoglobulin profile. Following a 24-hour stimulatory period, plasma/culture supernatants of each of the stimulatory conditions for all sample types was collected and the immunoglobulin profile determined using Luminex. The Immunoglobulin isotype levels depicted are reported as a percentage of the total immunoglobulin within a given sample. Median with interquartile range plotted. Statistical differences between culture conditions was calculated using a four-way mixed model ANOVA. Comparisons within groups was calculated using the Fishers LSD post-hoc test. A two-way step-up Benjamini, Krieger and Yekutieli False Discovery rate (FDR) approach, with a FDR of 1%, was used to correct for multiple testing. Statistical significance is indicated by an asterisk, in which the p < 0.05 (*), p<. 0.01 (**) and p<0.001(***). (a) Effects of sample type i.e. microenvironment complexity on immunoglobulin isotype abundance following antigenic stimulation (b) Effects of Quantiferon (QFN) status on immunoglobulin isotype abundance following antigenic stimulation (c) Effects of stimulatory conditions on immunoglobulin isotype abundance following antigenic stimulation

Additionally, the effects of QFN status and stimulation conditions on B-cell development was investigated. For majority of the investigated populations, QFN status was found to have no significant effect on the observed B-cell frequencies (Fig. 3B). A general trend of decreased frequencies for all populations was observed for QFN negative individuals compared to QFN positive individuals. In accordance, a significant decrease in the observed frequency of CD19^+^CD21^+^CD23^+^ (T2) B-cells (p=0.0012) was observed for QFN negative individuals. Furthermore, the effects of stimulation condition were examined (Fig. 3C). A common shift of increased B-cell frequencies was observed for most of the population subsets investigated following TLR9a stimulation compared to unstimulated controls. A significant increase in the frequency of CD19^+^CD21^+^CD23^−^ (MZ) B-cells was observed following TLR9a stimulation (p=0.0000) and *M.tb* infection (p=0.0000), compared to unstimulated cells. Similarly, a significant increase in CD19^+^CD21^+^CD23^+^ (T2) B-cells and CD19^+^CD21^−^CD23^+^ (FO) B-cells was observed in response to TLR9a stimulated compare to unstimulated (p=0.0000; p=0.0061) and H37Rv stimulated (p=0.0000; p=0.0198) samples, respectively. Contrariwise, a significant decrease in CD19^+^CD21^−^CD23^−^ (T1) B-cells was observed for samples stimulated with either TLR9a (p=0.0000) or H37Rv (p=0.0000) compared to unstimulated controls.

## Discussion

The standard application of isolated cell studies for the investigation of cell function have proven to be invaluable, however these studies do not account for the multi-faceted effects that surrounding cells types have on each other *in vivo*. This cellular communication results in physiological events that shape the immune response to various antigenic stimuli. In accordance, the purpose of this pilot study was to determine the effect of microenvironment complexity on B-cell function and to determine the significance of the use of isolated cell culture techniques in studies inferring the role of B-cells during TB disease.

Current research findings have signified the importance of B-cells during *M.tb* infection, in which absence or impaired function of this immune cell type has been associated with poor disease prognosis (Achkar et al., 2015; Bénard et al., 2018; Du Plessis et al., 2016a, 2016b; Rao et al., 2015). For decades, the primary function of B-cells was considered to be antibody secretion, forming part of the adaptive humoral response (Abbas et al., 2014; Zabriskie, 2009). These humoral immune responses were considered to be effective in controlling the growth and survival of extracellular invading pathogens exclusively. However, recent investigations analysing the efficiency of antibody-mediated immunity against several intracellular pathogens, including *M.tb*, have since disproven this notion (Achkar and Casadevall, 2013; Chan et al., 2014). In addition, studies have indicated non-humoral B-cell function, such as immune modulation through receptor engagement and cytokine expression, as key mechanisms by which these cells contribute to the successful control of *M.tb* infection (Bénard et al., 2018; Du Plessis et al., 2016b; van Rensburg and Loxton, 2018b). As such, the influence of *in vitro* isolated cell culture studies on B-cell development and function is of great importance, as currently observational findings inferring the physiological role of these cells during TB disease utilize these techniques (Du Plessis, Kleynhans, *et al.*, 2016; van Rensburg and Loxton, 2018b) and form the foundation upon which new TB drugs, host-directed therapies and TB vaccines are based.

In this study, antibody profiles were assumed to be a direct indication of the relative functional capacity of B-cells within the investigated samples. Notable, the presence of circulating antibody within the plasma samples compromises the inference of B-cell activity within whole blood samples and is a limitation of the study. Considerable changes in the immunoglobulin profile were observed across the different sample types, in which the relative percentage contribution of each of the measured isotypes, with the exception of IgG4, differed significantly (Fig. 1). Sample type, rather than stimulation condition, had a significant effect on the observed immunoglobulin profile. More specifically, a significant decrease in the relative abundant of IgG1 was observed following PBMC isolation compared to whole blood samples, while a significant increase in the relative abundance of IgG3 was found. The same pattern in the immunoglobulin expression was observed when comparing isolated B-cell samples with PBMCs. Importantly, the observed ‘increase/decrease’ in antibody levels is not equivalent to the concentration of these isotypes within a given sample but rather indicates the relative immunoglobulin diversity within the cellular microenvironment. The physiological implications of altered immunoglobulin production have been extensively reviewed in several disease states, where deficiency has been associated with increased susceptibility to bacterial infection (Franz et al., 1997; Hermans et al., 1976).

Immunoglobulins was shown to have a half-life of between 5-21 days (Anderson et al., 2006; Kim et al., 2007). Thus, circulating levels of antibodies were present within the plasma of whole blood samples prior to stimulation, whereas cells within the PBMC and isolated B-cell fraction were incubated in fresh media (Fig. 2A). This may have resulted in possible artefactual observations in the relative reduction in immunoglobulin levels for whole blood samples compared to PBMC and isolated B-cell samples. As such, significance of the observed [Ig] decrease was only considered between PBMCs and isolated B-cells. Collectively, these results illustrated that isolation procedures profoundly hindered the ability of B-cells to secrete several immunoglobulin isotypes; underscoring the fact that the presence of additional cell types is required for augmented B-cell activation and function.

Research has implicated IgA and IgG as leaders in protective anti-TB humoral immunity (Balu *et al.*, 2011; Achkar, Chan and Casadevall, 2015; Abebe *et al.*, 2018). The exact mechanisms by which these immunoglobulins achieve the protective effect is still unknown, and further investigation into their cellular targets is needed to better understand the role they play during TB disease (Li *et al.*, 2017).

Investigation of isolated cell studies on B-cells at the site of infection may provide valuable insight into anti-TB immunity. Mechanisms by which T-cells and dendritic cell induce this process are, directly through cell-cell interactions with adjacent B-cells and indirectly through the secretion of various soluble molecules (Fayette et al., 1997; Le Bon et al., 2001b). Studies utilising animal models to study anti-TB humoral immunity *in vivo* and *ex vivo* analysis of human samples from healthy and active TB participants has proved that humoral responses do in fact aid in the defence against *M.tb* infection (Achkar and Casadevall, 2013; Achkar et al., 2015; Chan et al., 2014). This further emphasizes the need to validate experimental observations in several independent experiments, utilizing difference techniques in health and disease.

The effect of sample type of B-cell development was investigated via evaluation of the relative frequencies on T1, T2, MZ and FO B-cells within each sample type following antigenic stimulation (Fig. 3). Interestingly, no significant difference in the frequency of all B-cell subsets was observed between whole blood and PBMCs. The involvement of MZ B-cells in T-independent early adaptive immune responses is well established, in which increased plasma cell differentiation and superior induction of Th1 expansion has been shown by MZ B-cells in comparison to FO B-cells (Attanavanich and Kearney, 2004; Lopes-Carvalho, Foote and Kearney, 2005; Crawford *et al.*, 2006). As such, MZ B-cells are regarded as primarily responsible for protective humoral and effector T-cell immune response. In contrast, activation of FO B-cells occurs via T-cell dependant mechanisms, and is thus involved in late immune responses (Martin, Oliver and Kearney, 2001; Balázs *et al.*, 2002). These results indicate the potential impact of cell isolation on B-cell derived immune responses; in which decreased frequencies of MZ B-cells was found. Thus, impaired B-cell development, as a result of diminished microenvironment complex due to cellular isolation, may result in the manifestation of inappropriate cellular responses to antigenic stimulation *in vitro*.

The effect of stimulation condition (TLR9 vs H37Rv) on immunoglobulin isotype abundance was investigated. A general pattern of enhanced B-cell development and increase of MZ B-cells was observed following TLR9a stimulation. The observed results are in agreement with previous findings indicates that *M.tb* challenge evokes early non-humoral B-cell responses that may play an importance role in the host defence against *M.tb* infection (Lenert *et al.*, 2005). These results underscore the ability of B-cell to actively respond to *M.tb* challenge and suggest that B-cells may be involved in directing and shaping the immune response during TB disease.

Our study demonstrates the influence that microenvironment complexity (i.e. sample type) had a profound impact on the activation and function of B-cells. These complex interactions underscore the basis for the use isolated cell studies to investigate cellular function, in an attempt to limit the degree of external factors influencing the observed results. This allows for the assumption, with complete certainty that the measured output for a cell population is in response to a particular drug or stimulus. However, it is important to remember that these isolated interactions are not indicative of whole blood scenarios (Brodin, Duffy and Quintana-Murci, 2019). Composite cellular interactions exist *in vivo* that may drastically influence the function of the (B-) cell type of interest resulting in a different reaction of these cells to the same drug or stimulus in whole organism.

## Methods

### Ethics Statement

Ethical approval was obtained from the health research ethics committee of Stellenbosch University (N16/05/070) and the City of Cape Town City Health. The study was conducted according to the Helsinki Declaration and International Conference of Harmonization guidelines. Written informed consent was obtained from all study participants.

### Study Participants

For this pilot study, we recruited 23 healthy individuals (15 individuals with a negative Quantiferon (QFN) status). The positive QFN status was suggestive of exposure to *M.tb*. Recruited participants did not present with any clinical symptoms of TB and had no previous record of active disease. All participants for this study were HIV negative. QFN positive and negative participants were matched according to socio-economic background.

### B-cell Isolation

Heparinized peripheral blood (18ml) was collected, of which 3mL was set aside for stimulation. From the remaining whole blood, peripheral blood mononuclear cells (PBMCs) were isolated using the ficoll-histopaque (GE Healthcare Life Sciences, USA) density gradient method. A fraction of the PBMC’s (3×10^6^ cells) was set aside for stimulation. Subsequently, B-cells were negatively isolated from the remaining PBMCs using MACS bead technology, according to the manufacturer’s instructions, with the B-cell isolation kit II (Miltenyi Biotec, South Africa). Once all the sample fractions had been collected, the cells were stimulated as described below. Purity of the enriched B-cell samples was confirmed by flow cytometry using anti-human CD19 mAb. All samples with resulting gated purity of above 90% were included in analysis.

### *In vitro* Stimulation assays

Whole blood (1mL blood/well), PBMC’s (1×10^6^ cells/well) and isolated B-cells (100 000 cells/well) were stimulated under 3 conditions: unstimulated, H37Rv (1×10^6^ CFU) or TLR9a (Sigma, USA) at 50 ng/ml. PBMCs and isolated B-cells were cultured in 96-well round-bottom plates in a total of 200uL complete media (RPMI plus L/Glutamine (Sigma, USA)) supplemented with 10% Fetal Calf Serum (FCS, Lonza, South Africa). Whole blood samples were incubated in a 24-well flat bottom plate in a total of 1.1mL (stimulants were diluted in complete media and added to 1mL blood). All sample fractions were incubated at 37°C and 5% CO_2_ for 24 hours. Following incubation, the plasma (in case of whole blood) and culture supernatants (in the case of PBMCs and isolated B-cells) were harvested, passed through a filter of 0.22μm (to remove any bacilli that may be contained within the sample) and stored at −80° for measurement of immunoglobulin secretion. The cells were then fixed with 4% paraformaldehyde (eBioscience, USA) for 30min at 37°C, washed with phosphate buffered saline (PBS (Lonza, South Africa) and cryopreserved (90% FCS and 10% Dimethyl sulfoxide (DMSO), Sigma-Aldrich, St, Louis, MO) in liquid nitrogen for future analysis by flow cytometry.

### Immunoglobulin isotype analysis by Luminex technology

Quantification of immunoglobulins within the plasma and culture supernatant, following the 24-hour stimulation, was determined using the MAGPix and Bioplex platforms (Bio-Rad Laboratories, California, USA). The immunoglobulins included IgA, IgE, IgG1, IgG2, IgG3, IgG4 and IgM. The experiments were performed according to the kit manufacturer’s recommendations.

### Phenotype Analysis by Flow Cytometry

The antibody panel for cell surface receptor analysis consisted of: CD19-BV605, CD21-PE/Dazzle, CD23-BV421, CD5-PerCP/Vio770, CD125 (IL5RA)-PE (All from Biolegend, California), CD3-FITC (BD, Germany) and CD178 (FASL)-APC (Miltenyi Biotec, South Africa). Cells were stained for 1 hour at room temperature in the dark, washed and acquired on a BD LSR II (BD Biosciences). The resulting data was analyzed using FlowJo v10 software (Treestar, USA).

### Statistical Analysis

Quantification of the immunoglobulin isotypes by Luminex was expressed as a percentage of the total immunoglobulin and statistical analysis performed on the relative percentage contribution of each of the isotypes within a sample. Data analysis for all Luminex results was performed using Statistica 12 software (Statsoft, Ohio, USA) and Prism 7 Software (San Diego, CA). Raw data was checked for normality using normality plots. Statistical differences between sample fractions (Whole Blood, PBMC, B-cells), QFN status and culture conditions (unstimulated, TLR9a, H37v) was calculated using a four-way mixed model ANOVA. Comparisons within groups was calculated using the Fishers LSD post-hoc test. A two-way step-up Benjamini, Krieger and Yekutieli False Discovery rate (FDR) approach, with a FDR of 1%, was used to correct for multiple testing. Data analysis of the flow cytometric plots was done using FlowJo V10 (Treestar, USA) and the resulting cell frequencies analyzed using Statistica and Prism 7 Software. Raw data was checked for normality using normality plots and winsorised using the Huber mean and MAD as necessary to reduce the residual distribution into close agreement with a normal distribution. Statistical significance is indicated by an asterisk, in which the p < 0.05 (*), p<. 0.01 (**) and p<0.001(***) or by letters, in which groups denoted with different letters indicate statistical differences.

## Acknowledgements

We thank the study participants for their participation and the Immunology Research Group laboratory where the assays were performed.

